# Invadopodia degrade ECM in the G1 phase of the cell cycle

**DOI:** 10.1101/412916

**Authors:** Battuya Bayarmagnai, Louisiane Perrin, Kamyar Esmaeili Pourfarhangi, Bojana Gligorijevic

## Abstract

Invadopodia are cancer cell protrusions rich in structural proteins (e.g. Tks5, cortactin) and proteases (e.g. MT1-MMP) and are responsible for degradation of the extracellular matrix (ECM). Tumor cell invasion and metastasis require cancer cells to be both proliferative and invasive, i.e. migrate through the tissue and assemble invadopodia. While several studies addressed how cell motility parameters change throughout the cell cycle, the relationship between invadopodia and cell cycle progression has not been elucidated. In this study, using invadopodia- and cell cycle- fluorescent markers, we show in 2D and 3D cell cultures, as well as *in vivo*, that breast carcinoma cells assemble invadopodia and invade into the surrounding ECM preferentially during the G1 phase of the cell cycle. Cells synchronized in the G0/G1 phase of the cell cycle degrade at significantly higher levels during the first 20 hours post-synchronization release. Consistent with this, mRNA and protein levels of the invadopodia key components, cortactin and MT1-MMP, peak at 14 hours post-release. Cell cycle progression is faster in cells in which invadopodia are abolished (by Tks5 knockdown), evidenced by earlier induction of cyclins A and B. A close look at the regulators of G1 revealed that the overexpression of p27^kip1^, but not p21^cip1^, causes faster turnover of invadopodia and increased ECM degradation. Furthermore, both endogenous and over-expressed p27^kip1^ localizes to the sites of invadopodia assembly. Taken together, these findings suggest that invadopodia function is tightly linked to cell cycle progression and is controlled by specific cell cycle regulators. Our results caution that this coordination between invasion and cell cycle must be considered when designing effective chemotherapies.

## Introduction

Metastasis is a complex, multi-step process that is initiated when cancer cells in the primary tumor acquire invasive properties, including motility and the ability to breakdown the extracellular matrix (ECM)^1^, and is responsible for the majority of cancer-related mortalities^2^. In breast cancer, the cellular structures responsible for ECM degradation are invadopodia. Invadopodia are small membrane protrusions, rich in actin and actin-binding proteins, such as cortactin and Tks5^3-8^. Upon maturation, matrix metalloproteinases (MMP2, MMP9, MT1-MMP) are delivered to the invadopodium, leading to the degradation of the surrounding matrix^9^.

We previously demonstrated that invasive cells oscillate between two distinct states, termed the Invadopodia state and the Migration state^10^. The Invadopodia state is characterized by cell stasis and the presence of invadopodia, whereas the Migration state is defined by cell translocation and the absence of invadopodia. This dynamic oscillatory behavior is regulated by the activity of ECM receptor integrin β1, which determines the length of time a cell spends in each state. Within the tumor microenvironment *in vivo*, invadopodia assembly is stimulated in areas rich in large blood vessels, macrophages and highly cross-linked ECM^11^. Perturbing any of these parameters has a direct effect on invadopodia function, establishing a strong *in vivo* correlation between the tumor microenvironment and invadopodia-driven motility. However, under these microenvironmental conditions, only 15% of tumor cells assemble invadopodia at any given time, while the rest are non-motile. This suggests the existence of additional intrinsic factors that determine the formation of invadopodia.

Deregulated cell cycle is a hallmark of cancer^12^. Cell cycle progression is governed by a complex network of cyclin-dependent kinases that define not only the phase of the cell cycle, but also the timing of transitions between phases^13^. In recent years, cell cycle regulators have been shown to exhibit roles in both tumor suppression and tumor promotion, particularly cyclin-dependent kinase inhibitors (CKI) p27^kip1^ and p21^cip114^. The mislocalization of these CKIs to the cytoplasm is associated with higher aggressiveness of several types of cancer, including breast cancer^15-20^. Most recently, a study demonstrated that p27^kip1^ facilitates invadopodia dynamics through the PAK1-cortactin axis^21^. Thus, further investigation is required to determine the molecular mechanism by which invadopodia-driven motility is coordinated with cell cycle progression.

Transcriptional profiling of invasive cells isolated from primary tumors revealed that in addition to the invasion gene signature, these cells exhibit a proliferation gene signature^22^. However, it is unclear whether cancer cells switch between invasive and proliferative phenotypes or display both phenotypes simultaneously. The Go or Grow hypothesis proposes that proliferation and motility are mutually exclusive. This dichotomy was tested in a number of cancer types, resulting in studies either supporting, e.g. in glioblastoma^23^, or challenging, e.g. mesothelioma and lung cancer^24^ this notion. This discrepancy could be attributed to differences in cancer types, or to the fact that the definition of “Go” largely focused on migration (i.e. translocation of the cell without breachment of ECM), and not invasion (i.e. motility involving an active breakdown of ECM). In melanoma, cancer cells undergo a phenomenon termed phenotype switching, where they switch from a highly proliferative and less invasive state to a highly invasive, but less proliferative state, thus, alternating between proliferation and invasion^25, 26^. This phenotypic plasticity in melanoma is deemed to be similar to the epithelial-to-mesenchymal (EMT) transition in other cancer types and key to developing resistance to chemotherapies^27^. Therefore, it is critically important to understand the relationship between proliferation and motility and distinguish between migration and invasion.

Towards determining how cell cycle progression is coordinated with cell migration in breast cancer cells, we recently introduced Fluorescent Ubiquitination Cell Cycle Indicator (FUCCI)-labeled cells into different microenvironments^28^. This study has shown that cell migration may occur during any of the cell cycle phases, except mitosis. In both 2D and 3D environments, speed and persistence of random migration did not vary between cells in G1 vs G2/M phases. However, in the presence of contact-guidance cues for directed migration, cells in G1 phase displayed more speed and persistence.

In this study, we investigate the coordination of cell cycle progression and invasion, specifically invadopodia. We demonstrate that during invasion, the Migration state may proceed without interruption through G1, S and G2 phases, whereas the Invadopodia state is cell cycle-dependent. The invadopodia function is enhanced in the G1 phase of the cell cycle. It is achieved by upregulating the gene and protein levels of invadopodia components MT1-MMP and cortactin, as well as by increasing the localization of Tks5 to invadopodia. Further, we show that matrix degradation and invasion are enhanced in cells synchronized in G1. This is true in 2D and 3D invasion assays and *in vivo*. We show that the CKI p27^kip1^, but not p21^cip1^, is recruited to the sites of invadopodia assembly and regulates invadopodia turnover. Thus, we demonstrate that invadopodia function, and therefore the metastatic potential of tumor cells, is tightly linked to cell cycle progression.

## Results

### 1. Cells degrade and invade ECM during the G1 phase of the cell cycle

To visualize both cell cycle progression and invadopodia formation in the breast carcinoma MDA-MB-231 cell line, we stably expressed the FUCCI reporters^29^, with the actin reporter mCerulean3-Lifeact7^30, 31^ (Fig. 1A). The FUCCI system consists of two fluorescent constructs, mKO2-hCdt1 (red) and mAG-hGem (green). Cdt1 expression peaks in G1 and drops to background levels during early S phase. Reciprocally, geminin level is the highest in late S, throughout G2 and in early M phases and sharply declines in late mitosis, during which no fluorescence is detected in the cell^29^. In early S phase, when both probes are expressed, the cell nuclei appear yellow. Time-lapse imaging of FUCCI reporters in our system confirmed the expected oscillatory expression patterns of mKO2-hCdt1 and mAG-hGem fluorescence (Fig. 1C).

**Figure 1.**
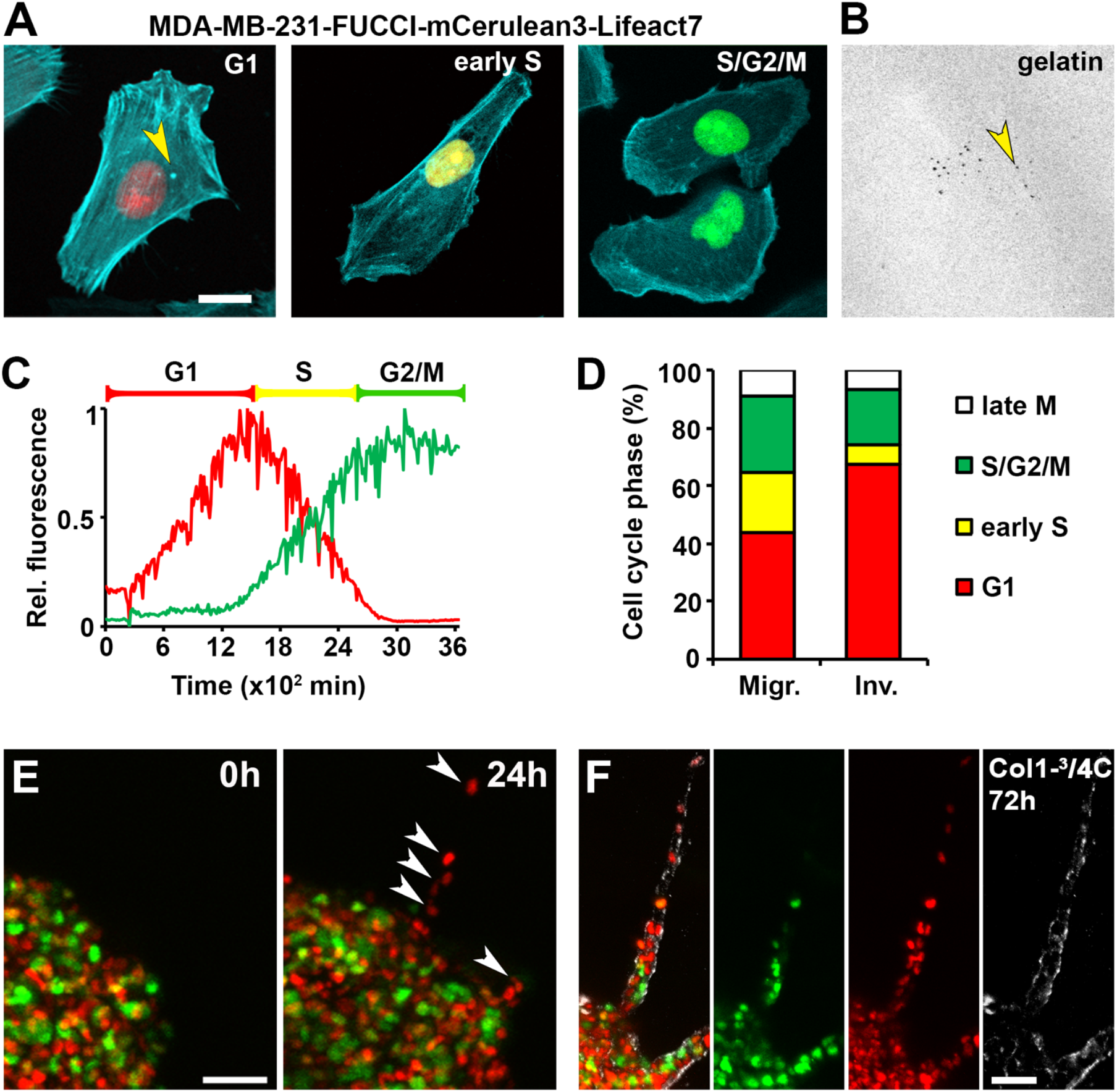
Invadopodia-directed ECM degradation occurs in the G1 phase of the cell cycle. A. Representative images of MDA-MB-231 cells expressing FUCCI cell cycle reporters and F-actin marker mCerulean3-Lifeact7 in G1 (left, invadopodia marked by yellow arrowhead), early S (middle) and S/G2/M (right) phases. Scale bar 10 µm. B. ECM degradation (degraded spot marked by yellow arrowhead) by invadopodia of the cell in A (left). C. Representative traces of red- and green- fluorescence of one MDA-MB-231-FUCCI cell over 3600 minutes. Brackets indicate how the cell cycle phases are classified based on the fluorescence signal: early S phase corresponds to the period where red fluorescence has a negative slope. The ratio of mean to maximum gray values is presented. D. Percentage of cells in each phase of the cell cycle in the Migration state (M, left bar) and the Invadopodia state (I, right bar) in an asynchronous population. Cells in the Migration state were defined by cell translocation and absence of invadopodia. Cells in the Invadopodia state were identified by the presence of actin puncta and degradation spots in the underlying gelatin layer. See Video S1, S2. E. Representative image of a 3D spheroid at 0h and 24h after embedding in high-density collagen I (5 mg/ml). Leading cells of the invasive strand are marked by white arrowheads. Scale bar 100 µm. See Video S3. F. Representative image of an invasive strand in a 3D spheroid at 72h after embedding in high-density collagen (5 mg/ml) merged (left) and individual channels showing FUCCI and degraded collagen antibody staining (Col1-¾C, right panel). Scale bar 50 µm.

In the G1 cells, we observed that actin was localized to small puncta characteristic of invadopodia precursors (Fig. 1A, left, yellow arrowhead), while puncta were absent from cells in early S and S/G2/M phases (Fig.1A, middle and right). This raised the question of whether the mature invadopodia and ECM degradation (Fig. 1B, yellow arrowhead) were also restricted to G1 phase of the cell cycle. To test this, cells were plated on fluorescent gelatin and simultaneously monitored for cell cycle, cell migration, presence of actin puncta and ECM degradation by live imaging over 40 hours. Interestingly, 68% of the cells in the Invadopodia state (mature invadopodia, plus ECM degradation) were in G1 phase, (Video S1, Fig. 1D), suggesting that cells engage in invadopodia-mediated ECM degradation preferentially during the G1 phase. From all cells in the Migration state^10^, 40% were in G1, 20% in early S phase and 30% in late S/G2/M phase, which mirrored the cell cycle distribution of the total population (Video S2, Figure1D).

To confirm our findings in a physiologically more relevant setting, we monitored the dynamics of MDA-MB-231-FUCCI-mCerulean3-Lifeact7 3D spheroid invasion into high-density collagen I, where cells utilize MMP-driven degradation of the matrix^32^ (Fig. 1E, F). In this assay, cells initiating invasion and strand formation were enriched for the G1 phase (Fig. 1E, white arrowheads, Video S3). Additionally, cells in the invasion strand actively engaged in degrading the surrounding matrix, which was confirmed by immunostaining for degraded collagen I (Fig. 1F). Collectively, these findings indicate that the MDA-MB-231 breast cancer cells degrade ECM and invade preferentially during the G1 phase of the cell cycle, both in 2D and 3D environments.

### 2. The expression and localization of invadopodia key components are cell cycle-regulated

Based on our findings so far, we reasoned that ECM degradation is more efficient in G1 phase compared to S/G2 phases of the cell cycle. To address this, we synchronized cells in early G1, expecting to see a robust increase in the amount of degradation compared to a cell population synchronized in S/G2 phase. Cells released from an early G1 arrest (synchronized by pre-treatment with lovastatin^33, 34^) showed a three-fold increase in ECM degradation compared to the asynchronous population (Fig. 2A_i_, 2A_ii_, 2A_iii_). In contrast, cells released from a S/G2 arrest (synchronized by pre-treatment with mitomycin C^35, 36^) showed a marked decrease in the ECM degradation (Fig. 2A_i_, 2A_iv_). These results demonstrate that ECM degradation is conducted at a higher efficiency during the G1 phase.

**Figure 2.**
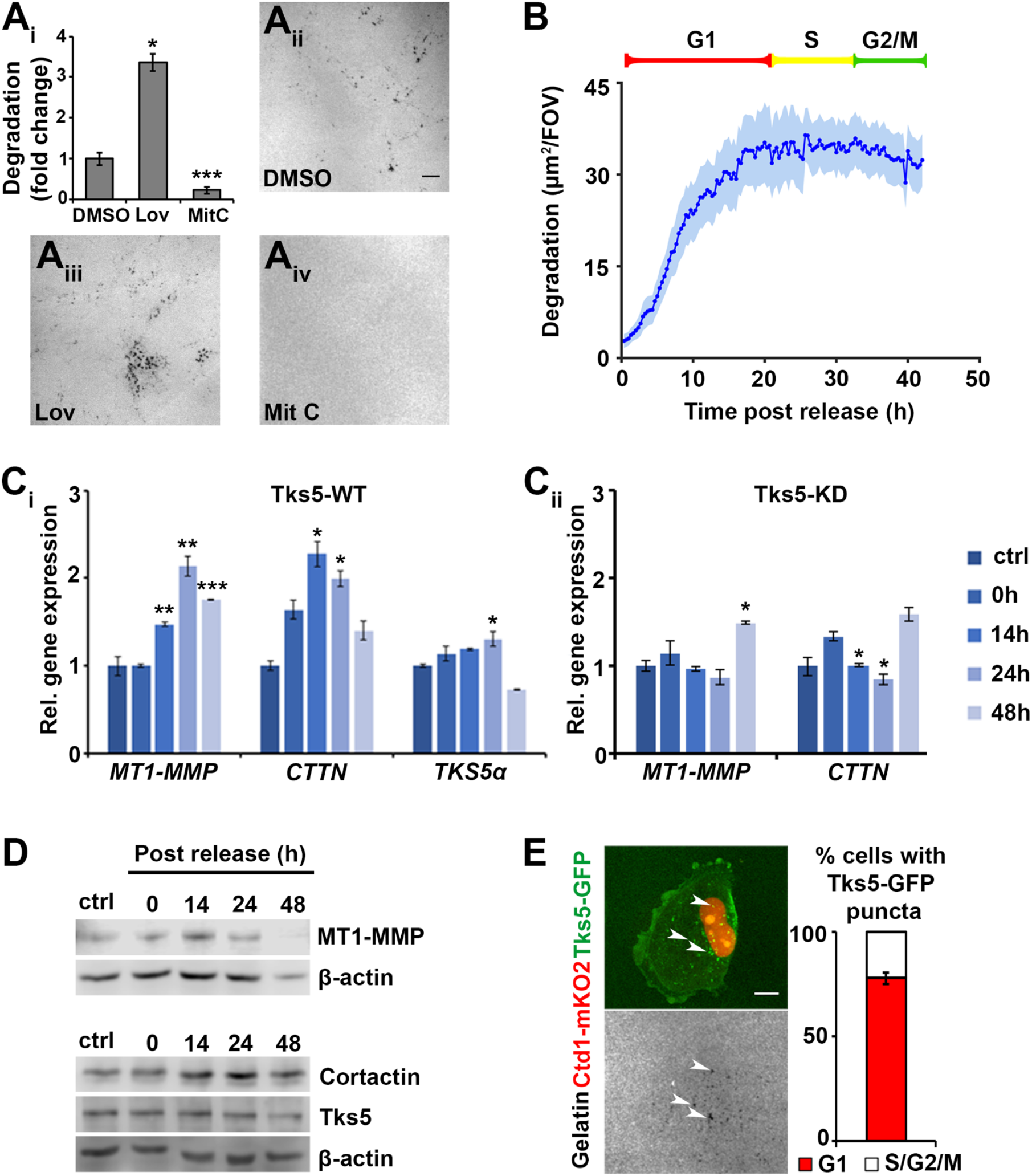
The invadopodia key components are cell cycle-regulated. A. Amount of degradation per cell in DMSO control and cells released from a synchrony in G1 (pre-treatment with 10µM lovastatin) or in S/G2 (pre-treatment with 1µg/ml mitomycin C). (A_i_) The results are shown as fold change relative to DMSO control. Representative images of degraded gelatin for DMSO control (A_ii_), lovastatin (A_iii_) and mitomycin C (A_iv_). *p<0.05, ***p<0.001 compared to DMSO control. B. Amount of degradation measured by time-lapse live imaging over 44 hours of cells released from G1 arrest (pre-treatment with 500nM PD0332991). The results are represented as a total degraded area per field of view. The shaded area in light blue indicates SEM. 5-10 cells/FOV, 10 FOV/replicate, 3 biological repeats were analyzed. Also see Video S4. C_i_. Gene expression levels of invadopodia key components, MT1-MMP, cortactin and Tks5, in cells that were harvested at 0-, 14-, 24-, and 48 h following release from the G1 arrest (pre-treatment with PD0332991). qPCR data were normalized to reference gene, *GAPDH*, and presented as fold change relative to the asynchronously cycling population. (C_ii_) Gene expression levels of invadopodia key components in the Tks5-KD cells. *p<0.05 compared to time 0h. D. Protein levels of invadopodia components of cells harvested from same conditions as in (C). β-actin is used as loading control. E. (Left) Representative immunofluorescence image of a G1 cell assembling Tks5-GFP-positive mature invadopodia (white arrowheads) and the underlying degraded gelatin (white arrowheads). (Right) Quantification of Tks5-GFP-positive puncta in G1 vs S/G2/M cells represented as the percent of total cells. Error bar represents SEM of two biological replicates, >50 cells/replicate.

In an asynchronously dividing population, ECM degradation exhibits a slow monotonic increase over time. If we monitor cells released from a G1 arrest, we expect to observe an exponential increase towards saturation in the amount of ECM degradation. To test this hypothesis, we monitored cells released from G1 synchrony using time-lapse imaging and measured the area of fluorescent ECM degraded over time. Consistent with our previous data, a sharp increase in the amount of degraded ECM occurred at 0-20 hours post-release from synchronization. The increase in degraded area plateaued at approximately 20 hours, indicating that the majority of degradation occurs during the G1 phase (Fig. 2B, Video S4).

Invadopodia assembly requires the orchestration of over 50 structural and enzymatic proteins^37^. We hypothesized that either the expression levels or the localization of core proteins will be enhanced to promote invadopodia in G1 phase. To test this, we measured the gene and protein expression levels of the invadopodia key components, including cortactin, Tks5 and the main metalloproteinase MT1-MMP, from cells harvested at 0, 14, 24 and 48 hours following release from G1 arrest. Consistent with the exponential phase of the ECM degradation, both gene and protein levels of cortactin, as well as MT1-MMP peaked at 14-20 hours (Fig. 2B-D). This dynamic gene expression disappeared when invadopodia were eliminated by Tks5 knockdown (Tks5-KD) (Fig. 2C_ii_).

Interestingly, Tks5 gene and protein levels remained relatively stable throughout the cell cycle, pointing to a different mechanism of regulation. Tks5 is heavily regulated at the level of recruitment and activity by phosphorylation of its tyrosine residues^38^. This prompted us to test whether its localization and recruitment to invadopodia were specific to the G1 phase. We transiently expressed the Tks5-GFP construct in MDA-MB-231 cells bearing only the Cdt1-mKO2 reporter (expressed in G1) and assessed the number of Tks5-GFP puncta localized to invadopodia in G1 vs S/G2/M cells. Consistent with the increased ECM degradation in G1, over 75% of cells containing Tks5-GFP-positive mature invadopodia were in G1 phase (Fig. 2E), indicating that Tks5 is recruited to invadopodia primarily during the G1 phase. Collectively, our findings suggest that invasion during the G1 phase is supported by an increase in the expression levels of the invadopodia key components cortactin and MT1-MMP and the localization of Tks5 to invadopodia.

### 3. Elimination of invadopodia accelerates cell cycle progression

Assembly of invadopodia is a structurally and metabolically intensive process, requiring a large number of actin regulators, proteases and kinases and both oxidative phosphorylation and glycolysis, suggesting high consumption of energym^37, 39, 40^. We reasoned that due to high energy demands and the assembly of multi-protein structures, cell cycle progression may be delayed in cells producing invadopodia. As Tks5 is an essential component of invadopodia assembly, we used the stable Tks5-KD cells to test the effect of invadopodia assembly on cell cycle progression.

First, we synchronized Tks5-WT and Tks5-KD cells in G1 phase. Next, cells were harvested at 4 hour intervals post-release. At each time-point, cell cycle phase distribution was assessed by propidium iodide (PI) flow cytometry and the induction of cyclin genes was measured using qRT-PCR. Expression of *CCNE1*, encoding for cyclin E, was induced 4 hours following release and persisted until 12 hours in both Tks5-WT and Tks5-KD cells, indicating that entry into S phase is timed similarly (data not shown). The expression of both cyclin A (*CCNA2*) and cyclin B (*CCNB1*) peaked 4 hours earlier in Tks5-KD cells compared to Tks5-WT, indicating that the progression through S and G2 phases occurs faster in the absence of invadopodia (Fig. 3A, 3B). Consistently, the PI flow cytometry results indicate that at 8-16 hours following release from G1 arrest the proportion of cells in S and G2 phases are higher in Tks5-KD cells (Fig. 3C, 3D). Cyclin D1 expression, encoded for by *CCND1* gene, remained stable throughout the cell cycle (data not shown), consistent with previous reports of cyclin D1 amplification in breast cancers^41^. Our findings suggest that in the presence of invadopodia MDA-MB-231 cells progress through the cell cycle slower, which demonstrates that there is a two-way interaction between invadopodia-driven invasion and cell cycle progression.

**Figure 3.**
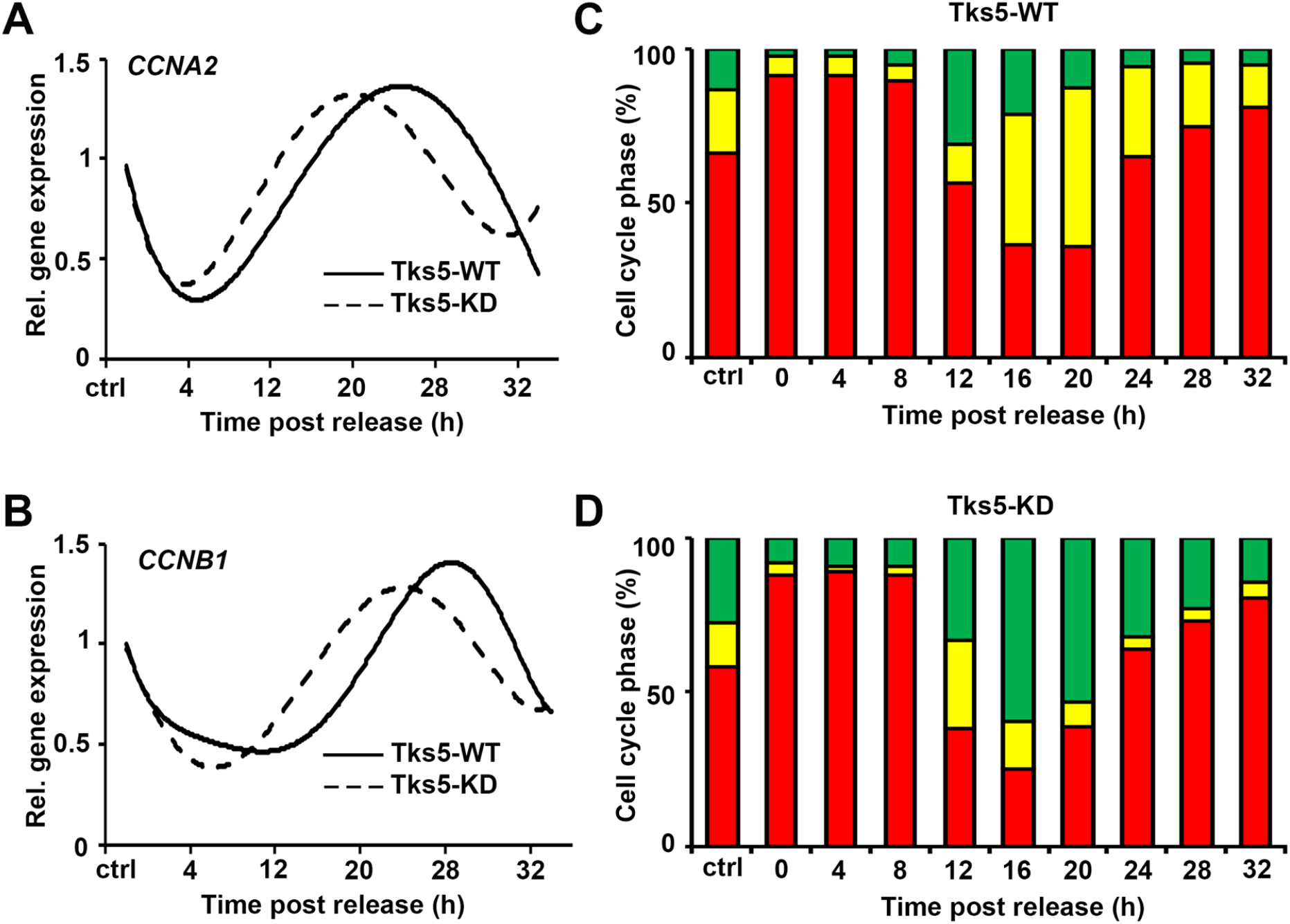
Elimination of invadopodia accelerates cell cycle progression. A and B. Gene expression level of cyclin A, *CCNA2* (A), and cyclin B, *CCNB1* (B), from Tks5-WT (solid line) and Tks5-KD (dashed line) cells harvested at 4h intervals post release from G1 arrest (pre-treatment with PD0332991). The qPCR data are normalized to reference gene, *GAPDH*, and presented as fold change relative to the asynchronously cycling population. Shown is the fitted trendline for the gene expression pattern. C and D. Cell cycle phase distribution of Tks5-WT (C) and Tks5-KD (D) cells harvested at 4h intervals post release from G1 arrest (pre-treatment with PD0332991) stained with PI and measured by flow cytometry. The 2N and 4N peaks were gated and quantified with FlowJo. The results are presented as percent of total cells measured at each time point.

### 4. Cyclin-dependent kinase inhibitor p27^kip1^ localizes to invadopodia and facilitates invadopodia turnover

The cyclin-dependent kinase inhibitors (CKI), p27^kip1^ and p21^cip1^, in addition to their roles in the cell cycle regulation, have also been implicated in pro-tumorigenic activities in the cytoplasm, such as inhibition of apoptosis^42^ and regulation of the cytoskeletal organization and cell migration^14^. Furthermore, the cytoplasmic pool of p27^kip1^ was recently shown to have a role in invadopodia turnover through the recruitment of PAK1^21^.

To gain insight into the molecular interplay that supports invadopodia function in G1 phase, we over-expressed p21^cip1^ or p27^kip1^ and assessed their effect on ECM degradation and invadopodia dynamics. Over-expression of p21^cip1^ resulted in no change in the amount of total degradation (Fig. 4A). Likewise, the number and size of degradation spots were similar compared to vector control (Fig. 4B, 4C). The size and the number of the degradation spots are known to reflect invadopodia lifetime^4^. Invadopodia exert mechanical forces onto the ECM via protrusion-retraction oscillations^10^. These oscillations, along with the delivery of MT1-MMP, lead to the degradation of the surrounding ECM, i.e. the “degradation spot” in 2D invadopodia assays. The number of degradation spots indicates the number of mature invadopodia assembled in a cell and the size of the individual degradation spots positively correlates with the number of protrusion-retraction cycles an invadopodium undergoes before its disassembly, and can be used as an indirect readout of invadopodia lifetime. In other words, the smaller the degradation spot, the shorter the lifetime and the higher the turnover^4^. Therefore, our results show that over-expression of p21^cip1^ has no effect on invadopodia function and thus is not involved in supporting invadopodia during G1.

**Figure 4.**
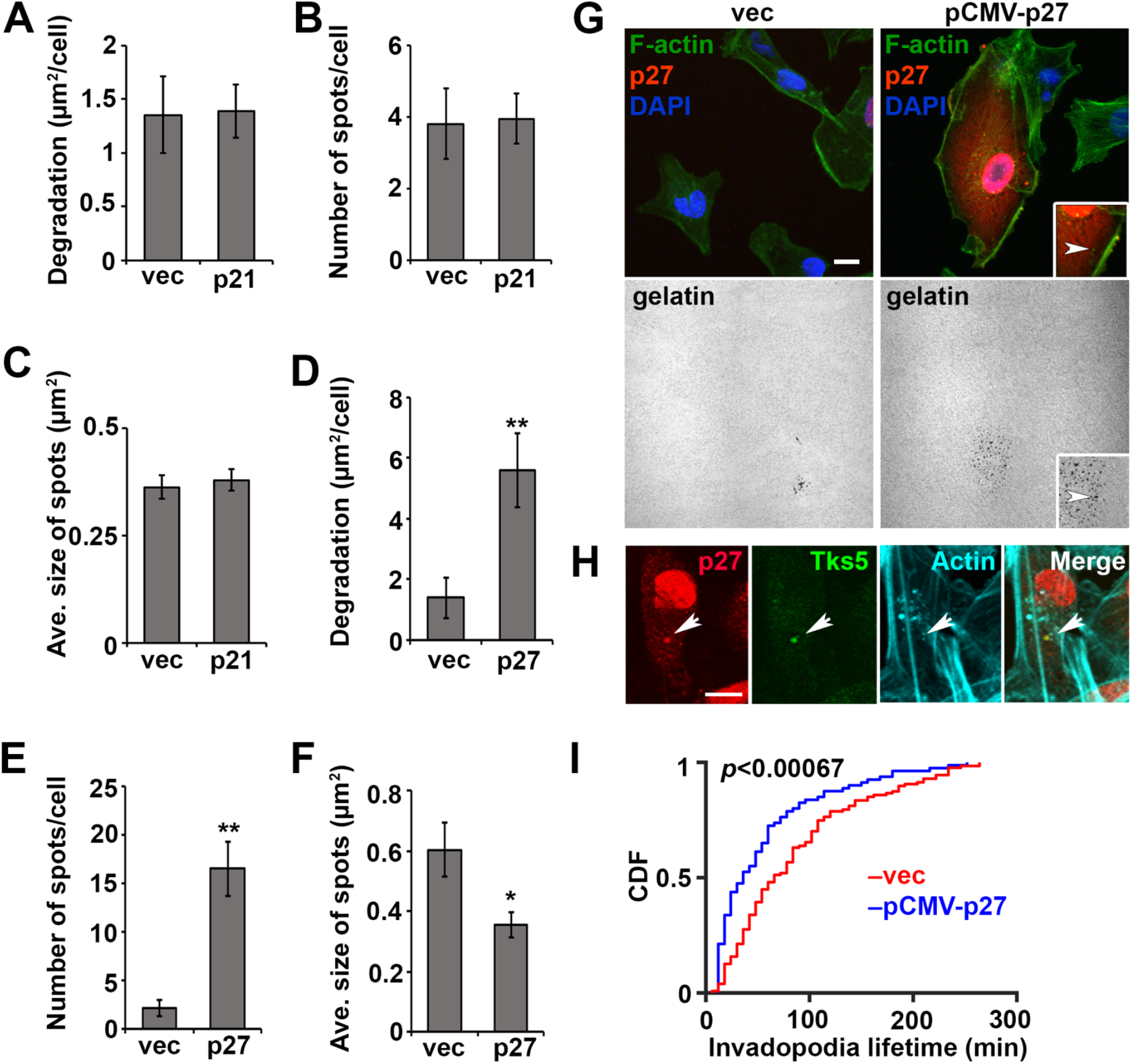
Cyclin-dependent kinase inhibitor, p27^kip1^, localizes to invadopodia and facilitates invadopodia turnover. Quantifications of the degradation area per cell (A), of the number of degradation spots per cell (B) and average size of the degradation spots (C) in vector control and cells over-expressing p21^cip1^. Error bars are SEM of three biological replicates. Quantifications of degradation area per cell (D), of the number of degradation spots per cell (E) and average size of the degradation spots (F) in vector control and cells over-expressing p27^kip1^. Error bars are SEM of three biological replicates. *p<0.05, **p<0.01 G. Representative immunofluorescence images of vector control and p27^kip1^ over-expressing cells with labeled nuclei (DAPI, blue), actin cytoskeleton (phalloidin, green), p27^kip1^ (red) and gelatin (gray). Insets are a zoomed-in area of p27^kip1^-overexpressing cell. White arrow points at the co-localization of actin, p27^kip1^ and the underlying degradation spot, indicating a mature invadopodium. Scale bar- 10µm. H. Immunofluorescence image of endogenous p27kip1 (red), Tks5 (green) and actin (cyan) in MDA-MB-231 cells. Co-localization of puncta is marked with white arrowheads. Scale bar 10 µm. I. Cumulative distribution function (CDF) plot of invadopodia lifetime measured from cells transiently transfected with vector control (red) or pCMV-p27^kip1^ (blue). Two biological replicates, >30 cells/replicate were analyzed. The distributions in these two conditions are significantly different with p<0.00067.

On the other hand, over-expression of p27^kip1^ led to a significant increase in the amount of degradation (Fig. 4D, 4G). Both over-expressed and endogenous p27^kip1^ was localized to the cytoplasm and recruited to the sites of invadopodia assembly, evidenced by its co-localization with actin and Tks5-rich puncta (Fig. 4G, 4H, white arrowheads). Furthermore, the increase in total degradation was due to an increase in the number of degradation spots (Fig. 4E) of smaller size (Fig. 4F), suggesting that p27^kip1^ not only stimulates invadopodia assembly, but also enhances invadopodia turnover suggested by smaller size of degradation spots. To test this, we performed live imaging on cells over-expressing p27^kip1^ and measured invadopodia lifetime by the stability of actin-rich puncta. In Fig. 4I, the values for lifetime were pooled and represented as a cumulative distribution function (CDF). Invadopodia with high expression of p27^kip1^ displayed significantly shorter lifetime (Fig. 4I). Collectively, our data indicate that p27^kip1^, but not p21^cip1^, is recruited to Tks5-positive invadopodia and facilitates invadopodia assembly and turnover.

### 5. Tumor cells *in vivo* form invadopodia in G1 phase

To determine whether the requirement for G1 phase is also true for invadopodia-driven invasion *in vivo*, we injected MDA-MB-231-FUCCI-mCerulean3-Lifeact7 cells orthotopically in the mammary fat pad of SCID mice. After 8-10 weeks, the xenograft tumors were exposed with a skin fold surgery and continuously imaged using intravital imaging with multiphoton microscopy^43, 44^. We recorded 4D stacks of tumor cells in the perivascular niche, where invadopodia phenotype is found, and assessed cell cycle phase of all cells and of cells forming invadopodia. We observed that the cells of each phase were evenly distributed throughout the tumor in any given field of view (Fig. 5A_i_). Next, we identified cells forming actin-rich protrusions, indicative of invadopodia (Fig 5A_ii_, Video S5) and quantified the number of G1 (red nuclei) vs G/M (green nuclei) cells forming such protrusions. We found that, in each field of view, G1 cells were more likely to show dynamic protrusions than G2/M cells (Fig. 5B). Thus, we demonstrated that in our model system, invadopodia *in vivo* are also assembled in G1 phase.

**Figure 5.**
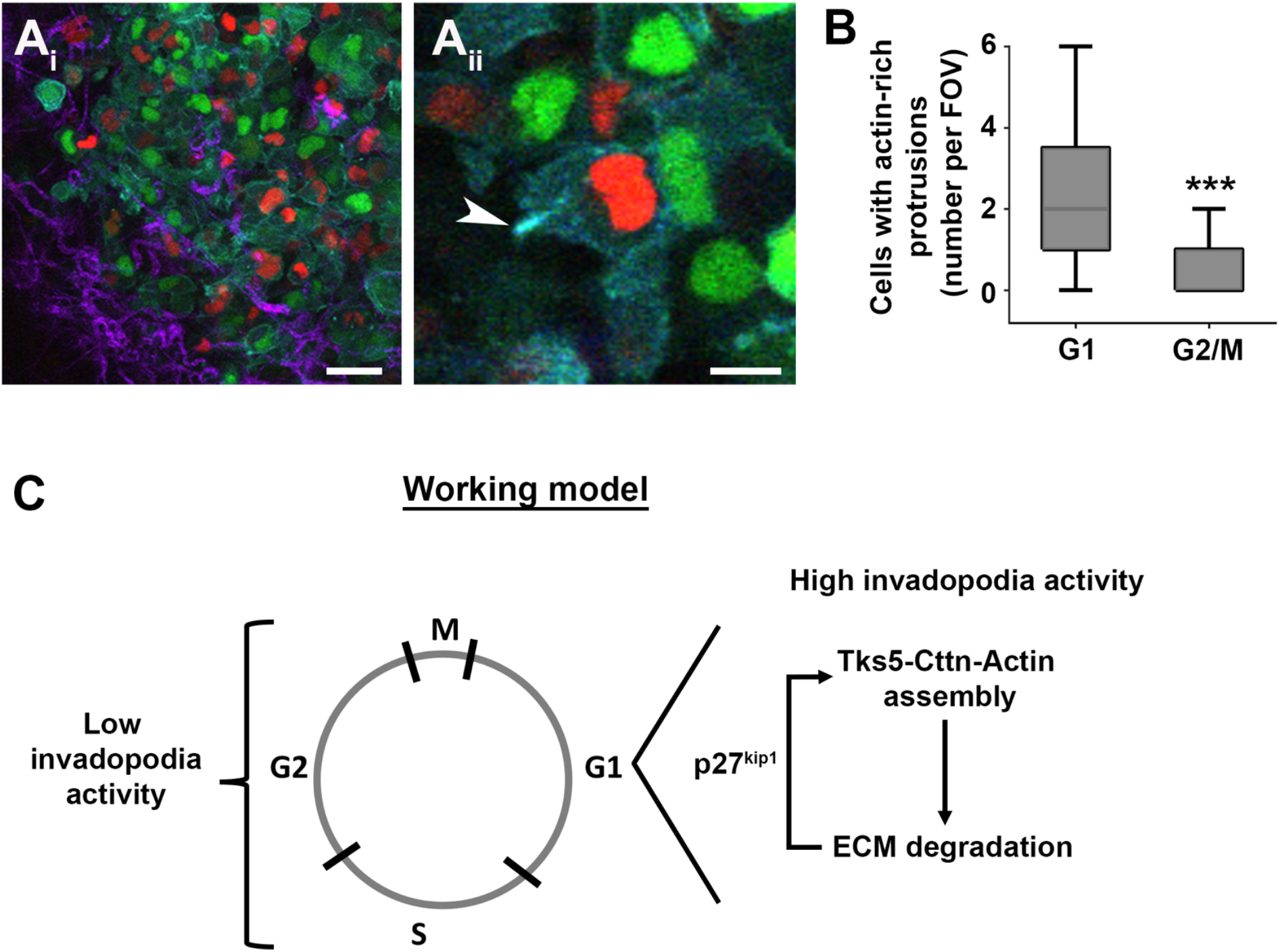
Invadopodia are enriched in G1 cells *in vivo*. A_i_. MDA-MB-231-FUCCI-Lifeact7-mCerulean3 cells in a mouse xenograft. Cells in G1 shown with red nuclei and S/G2/M in green, the actin cytoskeleton is in cyan and collagen fibers (SHG) are represented in purple. Scale bar- 50 µm. (A_ii_) Zoom-in on a cell in G1 phase (red nucleus) extending an actin-rich protrusion (arrowhead). Also see Video S5. Scale bar 10 µm. B. Quantification of cells forming actin-rich protrusions per phase of the cell cycle. The results are displayed as number of cells with actin-rich protrusions per field of view. >50 cells from 3 mice were analyzed. ***p-value<0.001. C. Proposed working model of the coordination of invadopodia function with cell cycle progression.

## Discussion

In this study, we show that invadopodia function is enhanced in the G1 phase of the cell cycle *in vitro* as well as *in vivo*. We demonstrate that the expression level of invadopodia key components (cortactin, MT1-MMP) is upregulated in G1 phase, while the localization to invadopodia of Tks5 is enriched in G1. The inhibition of invadopodia assembly by silencing Tks5 accelerates cell cycle progression. Investigation of the G1 regulators p27^kip1^ and p21^cip1^ revealed that the assembly and the turnover of invadopodia is mediated by p27^kip1^, but not p21^cip1^. Based on our findings, we propose a working model for the relationship between cell cycle progression and invadopodia as well as for the molecular interplay that leads to the enhancement of invasion in the G1 phase. We suggest that the cytoplasmic pool of p27^kip1^ in the G1 cells facilitates the assembly, maturation and turnover of invadopodia resulting in enhanced ECM degradation during G1 phase (Fig. 5C).

Our current study suggests that cell cycle status is another determinant of the Invadopodia state *in vivo*. Previously, we demonstrated that invadopodia assemble *in vivo* under specific extrinsic, microenvironmental conditions present in the perivascular niche and are necessary for metastasis^3, 11^. Interestingly, even in the perivascular niche, only 15% of cells exhibited invadopodia, raising the possibility that additional, intrinsic factors were required for invadopodia assembly. *In vivo* data gathered using MDA-MB-231-FUCCI-mCerulean3-Lifeact7 cells supports such a line of thinking, confirming that invadopodia are enriched in G1 cells.

The molecular relationship between cell cycle progression and invadopodia in cancer has not been studied. Recently, a mechanistic link implicating cell cycle regulators in invadopodia function has been revealed in *C. elegans*^45^. During development, the anchor cell (AC) requires an arrest in G1 for invadopodia-driven breakdown of the basement membrane. The G1 arrest is essential for invasion, as mutations leading to mitotic AC were shown to abolish invadopodia and invasion. This arrest is mediated by cyclin-dependent kinase inhibitor CKI-1 and histone deacetylase HDA-1, which promotes pro-invasive gene expression required for invadopodia formation and function in G1. The upregulation of pro-invasive genes is consistent with our findings that gene and protein levels of cortactin and MT1-MMP are increased in G1 in MDA-MB-231 breast cancer cells.

The role of the cytoplasmic pool of G1 CKIs, p27^kip1^ and p21^cip1^ in cell migration was previously established^14, 46, 47^. However, the relationship of these CKIs to invadopodia-driven motility has not been clarified. In a recent study, p27^kip1^ was shown to physically interact with cortactin and mediate its interaction with the p21-activated kinase, PAK1. PAK1 phosphorylates cortactin on S113 and thereby promotes invadopodia turnover^21^. In that work, the authors suggest that the role of p27^kip1^ in invadopodia function is restricted to the cytoplasmic compartment, without any links to its cell cycle-related function in the nucleus. Our data shows that the recruitment of p27^kip1^ to invadopodia and the invadopodia function are enriched in G1, suggesting that the cytoplasmic localization of p27^kip1^ may also be cell cycle-dependent. Hints supporting this come from a study done in Swiss 3T3 cells, which demonstrated that there is a substantial accumulation of cytoplasmic p27^kip1^ in G1 and that the ratio of the nuclear to the cytoplasmic pools changes with cell cycle progression^48^. Careful assessment of nuclear and cytoplasmic distributions of p27^kip1^ in cancer cells throughout cell cycle will help address this notion.

We showed that Tks5 localization to invadopodia is regulated by the cell cycle, while Tks5 mRNA and protein levels remain stable throughout cell cycle (Fig. 2E). In addition to controlling Tks5 localization, cell cycle may also be regulating its activation. Tks5 is heavily regulated by post-translational modifications, particularly by phosphorylation of its tyrosine residues. Recent large scale phosphoproteomics studies identified novel phospho-Serine/Threonine sites on Tks5, the phosphorylation of which varies depending on the cell cycle^49^. This finding is intriguing because key cell cycle regulators, such as CDKs, are Serine/Threonine kinases and thus potential modulators of Tks5 activity and recruitment. In fact, a member of the CDK family, CDK5, promotes invadopodia and invasion through the phosphorylation of actin regulator caldesmon^50^, warranting further studies on the effect of other CDKs on invadopodia function. Moreover, the acceleration of cell cycle progression in Tks5-KD cells suggests a reciprocal effect of invadopodia on cell cycle (Fig. 3). Given the potential of Tks5 as a prognostic marker of cancer progression in patients^51^, it is of utmost importance to further investigate the mechanistic link between Tks5 activity and cell cycle progression.

During development, G1 phase duration appears to dictate whether a stem cell remains proliferative or initiates a differentiation program^52, 53^. Namely, upregulation of cell cycle inhibitors, such as p27^kip1^ and Rb, which leads to lengthening of G1, promotes differentiation. Further supporting this notion, there appears to be a hierarchy in gene reactivation following mitotic exit. In hepatoma cells, genes required for the growth of the daughter cells are expressed first, followed by cell-cell adhesion genes consistent with the epithelial nature of the cell line. These genes precede the expression of liver cell type-specific genes suggesting that specialized genes are expressed last before the cell progresses to S phase^54^. This intimates that the emergence of new phenotypes may require lengthening of G1 phase for the full expression of the specialized genes. Given the fact that cancer cells often hijack developmental processes, it is tempting to speculate that the induction of the invadopodia behavior, accompanied by the high energy consumption of invadopodia assembly and function, may lengthen the duration of G1, which, in turn, may render cells insensitive to cytotoxic therapies.

Studies of tumor cell motility and metastasis are motivated by the need to develop effective therapeutics that can halt the spread of cancer cells to secondary locations, which is the main cause of cancer-related deaths. On the other hand, most clinically available chemotherapies target actively cycling cells^55^. It is becoming increasingly clear that both proliferation and motility characteristics of cancer cells need to be taken into account when designing therapies. We previously showed a direct link between invadopodia *in vivo* and metastasis^11^. Our current study indicates that breast carcinoma cells in G1 possess the highest capabilities for invadopodia function, and hence, metastasis. This suggests that anti-proliferative drugs may select for invasive cells and conversely, that targeting motile cells may promote the proliferative state and the growth of metastases. Taken together, our data suggests that to effectively target cancer metastasis, understanding the mechanistic relationship between cell cycle progression and invadopodia is essential.

## Materials and Methods

### Ethics statement

All experiments on animals were conducted in accordance with the NIH regulations and approved by the Temple University IACUC protocol number 4397.

### Cell lines and culture conditions

The cell cultures were maintained in DMEM media supplemented with 10% fetal bovine serum (FBS, Atlanta Biologicals) and 50 U penicillin/50 µg streptomycin (Corning) per ml.

MDA-MB-231-FUCCI stable cell line was generated by transducing FUCCI constructs, mKO2-mCdt1 (30/120) and mAG-hGem (1/110), with the replication-defective, self-inactivating lentiviral expression system. To visualize cell motility, the plasmid bearing actin-binding protein mCerulean3- Lifeact7 was introduced to MDA-MB-231-FUCCI cells by electroporation (Lonza) and selected for with 500 µg/ml geneticin. We performed fluorescence-activated cell sorting (FACS) after 14-20 days of selection and collected top 20% highest fluorescing cells (after excluding the top 5% highest fluorescing population). The mCerulean3-Lifeact7 expression was maintained under 500 µg/ml geneticin (Invitrogen). mCerulean3-Lifeact7 plasmid was a gift from Michael Davidson (Addgene plasmid #54721). FUCCI constructs were obtained from RIKEN Tsukuba BioResource Center (Japan). pCMV5 human p27^56^ was a gift from Joan Massague (Addgene plasmid #14049). Flag-p21 WT^15^ was a gift from Mien-Chie Hung (Addgene plasmid #16240). Tks5-GFP construct is a gift from Sara Courtneidge (OHSU).

Tks5KD cell line was generated by transducing MDA-MB-231-FUCCI-mCerulean3-Lifeact7 cells with lentiviral particles (5 particles/cell) containing shRNA in pLKO.1 vector targeting Tks5 (Sigma Aldrich MISSION library). Single cell colonies were selected with 0.5 µg/ml puromycin.

### 2D matrix degradation assay and immunofluorescence

35mm glass-bottom MatTek (MatTek Corporation) were coated with fluorescently labeled gelatin as previously described^57^. For cell cycle synchronization experiments, cells pre-treated with DMSO control, 10µM lovastatin (G1 phase) (PHR1285, Sigma-Aldrich) or 1µg/ml mitomycin C (S phase) (11435, Cayman Chemical) for 24 hours, then washed with PBS, trypsinized and re-plated on gelatin-coated glass-bottom dishes for 16-20 hours. The cells were fixed for 15 min with 4% paraformaldehyde, permeabilized for 5 min in 0.1% Triton X-100, blocked for 2 hours in 1% BSA /1% FBS in PBS, and incubated with primary antibodies for 1-3 hours, then with secondary antibodies, DAPI and phalloidin-633 (1:250) for 1-2 hours. Rabbit anti-p27^kip1^ (Abcam, ab32034, 1:100) and mouse anti-Tks5 (clone 13H6.3, MABT336, Millipore) were used. The samples were imaged in the laser scanning confocal mode of the hybrid confocal-multiphoton microscope (Olympus FV1200). Matrix degradation was quantified by thresholding the signal in the gelatin channel and measuring the number, size and the total area of degradation spots using the Particle Analysis tool in FIJI.

### 3D spheroid invasion assay

3D spheroids were generated by the hanging drop method. 5000 cells per 20µl drop containing 4.8 mg/ml methylcellulose, 20 µg/ml Nutragen (Advanced Biomatrix), were placed on the lid of tissue culture dishes. The lids were carefully turned and placed on the bottom reservoir of the dishes filled with PBS to prevent evaporation. The spheroids were allowed to form for 48 hours. Alternatively, spheroids were generated in a 96-well flat bottom dish, coated with 1.5% w/v agarose. Briefly, 0.3 g of agarose was added to 20 ml DMEM and autoclaved for 20 min at 120°C, 2 bar. 50 µl of 1.5% agarose was distributed to each well of a flat bottom 96-well plate and cooled to room temperature for approximately 20 min. The hermetically sealed plates can be stored for up to 10 days. The cells were trypsinized, counted and diluted to 2.5×10^4^ cells/ml in ice-cold medium. Matrigel (thawed on ice overnight) was added to final concentration of 2.5%. Cell mixture was distributed to the agarose-coated 96-well plate, with each well containing 5000 cells in 200 µl. All reagents were kept on ice and pre-chilled pipette tips were used. The plate was centrifuged at 4°C for 10 min at 1000 x *g* and spheroids were allowed to form over 72 hours at 37°C.

The fully formed spheroids were embedded in 50 µl of 5 mg/ml rat-tail collagen I gel (Corning, alternate gelation procedure was followed from manufacturer’s guidelines) and distributed to each well of a 24-well plate. Invasion was recorded by live imaging at 1 hour intervals over 70 hours, in the laser scanning confocal mode of the hybrid confocal-multiphoton microscope (Olympus FV1200). For immunofluorescence, the spheroids were fixed in 4% paraformaldehyde for 20 min, permeabilized in 0.1% TritonX-100 for 20 min, blocked in 1% FBS/1% BSA in PBS overnight at 4°C, incubated with primary antibody against degraded collagen I (Col1-C^3^/_4_, ImmunoGlobe, 1:100) overnight at 4°C and with secondary antibody for 3-4 hours at room temperature.

### 2D live cell imaging

5×10^4^ cells were seeded on gelatin-coated glass-bottom dishes and allowed to adhere for 2 hours before imaging. Live cell imaging was performed on a wide-field Olympus IX-81 microscope, equipped with an environmental chamber (In Vivo Scientific). The images were acquired at a single focal plane using an Olympus 60X oil immersion objective at CFP, FITC, TRITC, Cy5 and DIC channels. Images were acquired at 20 min intervals for 48-60 hours. For FUCCI expression validation, nuclei of single cells were tracked in both red and green channels and their mean gray values were recorded. For synchronization, cells were pre-treated with 500 nM PD0332991(PZ0199, Sigma-Aldrich) for 24 hours, washed with PBS and re-plated on gelatin-coated glass-bottom dishes before live imaging.

### Intravital imaging

1×10^6^ of MDA-MB-231-FUCCI-mCerulean3-Lifeact7 cells was suspended in 100 µl of 20% collagen I in PBS and injected orthotopically into the mammary fat pad of 5-7 week-old female SCID mice. After 8-10 weeks, when the tumor diameter reached 8-10 mm, the animals were surgically prepared for continuous intravital imaging. The tumor tissue was exposed by skin flap non-survival surgery as previously described^11, 43^. Briefly, anesthesia in mice was induced with 5% isoflurane, the levels of which were gradually decreased to, and maintained at 1.5%. The mice were kept in 37°C environmental chamber (In Vivo Scientific) and hydrated by intraperitoneal injection of 100 µl of warm, sterile PBS every hour.

4D stacks were collected in the multiphoton imaging mode of the hybrid confocal-multiphoton microscope (Olympus FV1200MPE), with 30X objective with silicone oil immersion (UPLSAPO 30X, NA 1.05, Olympus). The areas were selected based on the presence of flowing macrovessels, visible through the ocular using the green FITC filter, as the lack of green autofluorescence. While blood flow indicates proper circulation, large capillary diameter is indicative of the perivascular niches amenable for invadopodia assembly^11^. 4D stacks were collected at 3 min intervals for up to 6 hours. At the end of each imaging session, the animals were euthanized and the tumor sample was collected.

The acquired time-lapse videos were analyzed using FIJI. The frames were corrected for motion artifacts using HyperStackReg, an extension based on StackReg plug-in^58^. The cells extending invadopodia were identified based on the morphodynamic properties and scored for cell cycle phase based on the FUCCI signal.

### Gene expression qRT-PCR

Total mRNA was extracted with Trizol (Invitrogen) according to the manufacturer’s recommendations. The RNA yield was measured with NanoDrop (ThermoFisher). 1µg of RNA was used for Reverse Transcriptase PCR (High-Capacity cDNA Reverse Transcription Kit, Applied Biosystems) reactions. 1:20 dilution of cDNA was used for quantitative Real Time PCR (PowerUp SYBR Green Mastermix, Applied Biosystems). For Tks5, the primers detecting the most abundantly expressed isoform in adult tissues, *TKS5α*, were used^59, 60^.

### Western blot analysis

The cells grown on a gelatin layer were harvested in RIPA lysis buffer (cat. R3792, Teknova), supplemented with protease inhibitors (Roche, complete cocktail) and phosphatase inhibitors (Sigma, Halt cocktail). SDS-PAGE was performed with 20µg total protein per sample, transferred to a PVDF membrane (Immobilon), blocked with 5% BSA/TBST for 1 hour at room temperature and incubated with the primary antibody in 5% BSA/TBST overnight at 4°C. Rabbit anti-Tks5 (anti-SH3PXD2A, ProteinTech 18976-1-AP, 1:1000), mouse anti-Cortactin (Abcam, ab33333, 1:1000), rabbit anti-MT1-MMP (Cell Signaling, D1E4 13130S, 1:1000), and mouse anti-β-actin as loading control (Santa Cruz C4, sc-47778, 1:500) were used. The membranes were then incubated with the secondary antibody (1:5000) in 5% non-fat milk/TBST for 1 hour at room temperature. Bands were visualized using chemiluminescence detection reagents (WesternBright, Advansta) and blot scanner (C-DiGit, LI-COR).

### Cell cycle analysis with propidium iodide (PI) staining and flow cytometry

Cells of indicated treatments were cultured in 6-well dishes, harvested and re-suspended in ice-cold PBS containing 2% FBS. Cells were counted, spun at 1200 rpm for 5 min and re-suspended in 250 µl PBS. Cells were fixed by a drop-wise addition of 750 µl of ice-cold 95% ethanol, while vortexing gently at low speed and then incubated at 4°C overnight or up to one week. Cells were allowed to equilibrate to room temperature (~30 min) to loosen the cell pellet and then centrifuged at 2000 rpm for 5 min. The supernatant was aspirated and the cell pellet was re-suspended in 150 µl of the PI staining solution (50 µg/ml PI, 0.2 mg/ml RNase A, 0.02% Triton X-100, PBS) and incubated for 30 min at room temperature, protected from light. After the staining incubation, the samples were diluted with 150 µl of PBS in 10 mM EDTA to prevent clumps, and filtered through 40 µm mesh cell strainer. The samples were measured using BD Accuri C6 flow cytometer (BD Biosciences).

### Invadopodia lifetime assay

Time-lapse images of MDA-MB-231-mCerulean3-Lifeact7 cells transiently transfected with vector control and pCMV-p27 plasmids were acquired every 6 min for 24 hours and analyzed using FIJI. The invadopodia lifetime was defined as the time between the appearance and the disappearance of actin puncta. Actin puncta present throughout the entire stack were excluded from the analysis. The lifetime measurements are presented as Cumulative Distribution Function (CDF) and compared using Kolmogorov-Smirnov test.

## Acknowledgments

We are grateful for the Tks5-GFP construct and experimental advice provided by Sara Courtneidge and her group (OHSU). We are also grateful for the meaningful discussions and help with experimental design provided by Xavier Grana (Temple School of Medicine). This study was partially funded by R00 CA172360, National Institutes of Health and Concern Foundation Conquer Cancer Now Young Investigator Award to B.G.

## List of Abbreviations

2D: Two-dimensional
3D: Three-dimensional
4D: Four-dimensional
BSA: Bovine serum albumin
CDK: Cyclin-dependent kinase
CKI: Cyclin-dependent kinase inhibitor
CMV: Cytomegalovirus
DAPI: 4’,6-diamidino-2-phenylindole
DMSO: Dimethyl sulfoxide
ECM: Extracellular matrix
EDTA: Ethylenediaminetetraacetic acid
FACS: Fluorescence-activated cell sorting
FBS: Fetal bovine serum
FUCCI: Fluorescent Ubiquitination Cell Cycle Indicator Gem Geminin
KD: Knockdown
Lov: Lovastatin mAG mAzami Green
Mit C: Mitomycin C
mKO2: mKusabira Orange 2
PAK1: p21-activated kinase
PBS: Phosphate-buffered saline
PI: Propidium Iodide
qRT-PCR: Quantitative real-time polymerase chain reaction
Rb: Retinoblastoma
RT-PCR: Reverse transcriptase polymerase chain reaction
SCID: Severe combined immunodefficiency
SDS-: Sodium dodecyl sulfate- polyacrylamide gel
PAGE: electrophoresis
Ser: Serine
TBST: Tris-buffered saline- Tween 20
Thr: Threonine
Tyr: Tyrosine
WT: Wild type

